# Radiotherapy induces specific miRNA expression profiles in glioblastoma exosomes

**DOI:** 10.1101/2021.09.08.459416

**Authors:** Axel Boukredine, Sofiane Saada, Stéphanie Durand, Alexandre Nivet, Barbara Bessette, Amel Rehailia, Pierre Clavère, Marie-Odile Jauberteau, Elise Deluche, Fabrice Lalloué

## Abstract

**Background:** Glioblastoma (GBM) is the most aggressive and frequent primary brain tumor during adulthood. One of the major treatments is the association of surgery and a combination of chemo and radiotherapies. Despite its immediate efficiency, it fails to prevent the cancer recurrence in the irradiated area due to radioresistance mechanisms.

MicroRNAs (miRNAs or miR) are small non-coding, single strand RNA molecules encoding to various specific genes and able to regulate their expression and induce the tumor cell survival leading to radioresistance. Small extracellular vesicles (EVs), or exosomes released by tumor cells in tumor microenvironment and blood circulation are able to transport and diffuse miRNAs and affect the microenvironment by spreading the miRNAs, which drive radioresistance.

**Aims:**

i. To identify the variations of miRNAs expression induced by irradiation in human glioblastoma U87-MG cells and their secreted exosomes collected in supernatants.
ii. To analyze the miRNAs variations in EVs-derived from the plasma of patients during radiotherapy, in order to identify a miRNA signature induced by radiotherapy in a liquid biopsy.

**Materiel and methods:** U87-MG cells were cultured on plates and exposed to irradiation. miRNAs analyzes were performed in cells and in EVs isolated from cell supernatants to determine miRNAs expressions both in cells and in secreted exosomes before and after irradiation.

Plasma-derived EVs were collected from 4 glioblastoma patients before and after surgery and radiotherapy treatments.

**Conclusion:** The analysis of miRNAs expression profiles in both GBM cells and their derived EVs revealed that miR profile changes after irradiation. However, the number of similar miR between cells or EVs, following cell irradiation, was restricted to 3 miRs alone suggesting that the irradiation-induced changes in the miR profile in the cells and their EVs are not closely linked. In this context, the miR profile in EVs from patients plasma was investigated to establish a potential link with the miRNAs profile observed in EVs from irradiated cells and to assess its relationship with the response to radiotherapy. Three miRs (different from those identified in cells) were common between EVs derived from cells and patients derived-exosomes. These miRs detected in circulating EVs could provide a specific and reliable signature in response to ionizing radiation, which could be useful for monitoring the effectiveness of radiotherapy. Further experiments on a larger patients population with clinical data could also help to define whether this signature might have a prognostic value on the response to radiotherapy.

## Introduction

Glioblastoma is one of the most aggressive tumors with a median survival of 15 months and a particularly poor prognosis that has remained unchanged for more than 10 years. Following surgical resection, patients still treated with a combination of radiotherapy and chemotherapy. However, this therapeutic strategy often fails due in particular to the radio- and chemoresistance properties of tumor cells and the crosstalk between tumor cells and the microenvironment. Soluble factors often considered as the main communication mechanism between the tumor and its microenvironment. However, many biological interactions such as intercellular signaling or communication with extracellular vesicles could be decisive for tumor progression and the development of resistance mechanisms. The function and involvement of extracellular vesicles in resistance mechanisms are increasingly discussed and show changes in the content of exosomes derived from tumor cells in response to chemotherapy and ionizing radiations, suggesting that drug resistance could be mediated by EVs [1,2]. Since miR and mRNA were the first classes of nucleic acids characterized in exosomes, miRNAs derived from tumor exosomes are considered as mediators of tumor progression and metastasis [3–5]. Indeed, the horizontal transfer of exosomal miRNAs from cancer cells might influence and control its microenvironment by enhancing the tumor angiogenesis to promote their metastatic initiation [6]. Similarly, the transfer of genetic materials via exosomes into glioblastoma allows tumor cells to remodel their microenvironment in order to promote radio- or chemoresistance [7]. miRNAs profile of tumor cells are sensitive and modified according to the treatment. Thus, when tumor cells are exposed to stress induced by radiotherapy, their miRNAs profile changes depending on the treatment. In a similar manner, the miRNAs content of exosome released by cancer cells reflect the original miRNAs profile from these tumor cells. Differential exosomal miRNAs screening at different levels could be useful to reveal tumor aggressiveness and changes of response to radiotherapy. Progress in miRNAs profiling permit to analyze miRNAs content of the exosomal cargo and to compare it before and after radiotherapy. First evidence demonstrated that exosome release increase following ionizing radiations in different glioblastoma cell lines [8]. They could be involved in the activation of U87MG recipient cell migration. In our study, miRNA profile changes were observed in patients during Stupp protocol at 20 Gy and 60 Gy for analyzing whether stress induced by ionizing radiations is able to modify miRNAs profile. Although tumors have a different sensitivity to ionizing radiations, the high proliferative rate of glioblastoma cells confers them a highly sensitivity to radiations which explains why their proliferation is temporarily stopped after radiotherapy. However, the acquisition of radioresistance during treatment may result in local recurrence. The acquisition of radioresistance properties could rely on miRNA profile changes. That is the reason why in our study, we chose to determine whether stress induced by ionizing radiations is able to modify miRNAs profile in patients during Stupp protocol.

Therefore, we provide evidence that irradiation induces changes of miRNAs profile both in EVs secreted by irradiated glioblastoma cells and in plasma-derived EVs from glioblastoma patients during radiotherapy. We identified 4 miRNAs overexpressed upon radiotherapy that could be essential to regulate a common target already considered as a tumor suppressor gene. Altogether, our results suggest that the miRNAs composition in secreted microvesicles could reflect molecular changes in cells from which they derived and, therefore, may provide diagnostic information and a specific and reliable signature in response to ionizing radiation that could be useful for monitoring radiotherapy efficiency. Further experiments will have to be consider to determine the prognostic value of this new miRNAs profile on radiotherapy response from a larger patient population with complete clinical data.

## Materials and Methods

### Cell lines

The human GBM cell line U87-MG purchased from American Type Culture Collection (ATCC^®^). Cells were grown in complete medium containing Minimum Essential Medium (MEM) (Lonza^®^) supplemented with 10% FBS (Life Technologies^®^), 2% sodium bicarbonate, 1% sodium pyruvate, 1% non-essential amino acids solution and 1% penicillin/streptomycin at 37 °C in a humidified atmosphere of 5% CO2 and 95% air. For exosomes experiments, cells were cultured using exosome-free complete medium (EFM). The EFM obtained using exosome-depleted FCS after 16 h ultracentrifugation at 120,000 g.

### Cell line Irradiation

U87-MG cells were seeded in 6-well plates for 24 to 48 hours after sowing, transported to the radiotherapy department of the Limoges University Hospital. Cells were treated by radiotherapy using a plexiglas plate designed in the laboratory. The plate contains 8 radiotherapy catheters connected high flow iridium 192 source projector pass, which bring radiotherapy source directly to cells and deliver (Figure 1 a). The established predictive dosimetry showed good coverage of the targeted cells (Figure 1 b). The irradiation delivered a dose of 7 Gy, in 20 minutes irradiation. Irradiation process were assessed in a radioprotection bunker, under the supervision of the staff of the Radiotherapy department Limoges University Hospital. Cell were brought back to the laboratory and returned to incubation and cultured for 48 h under previously described conditions.

**Figure 1:**
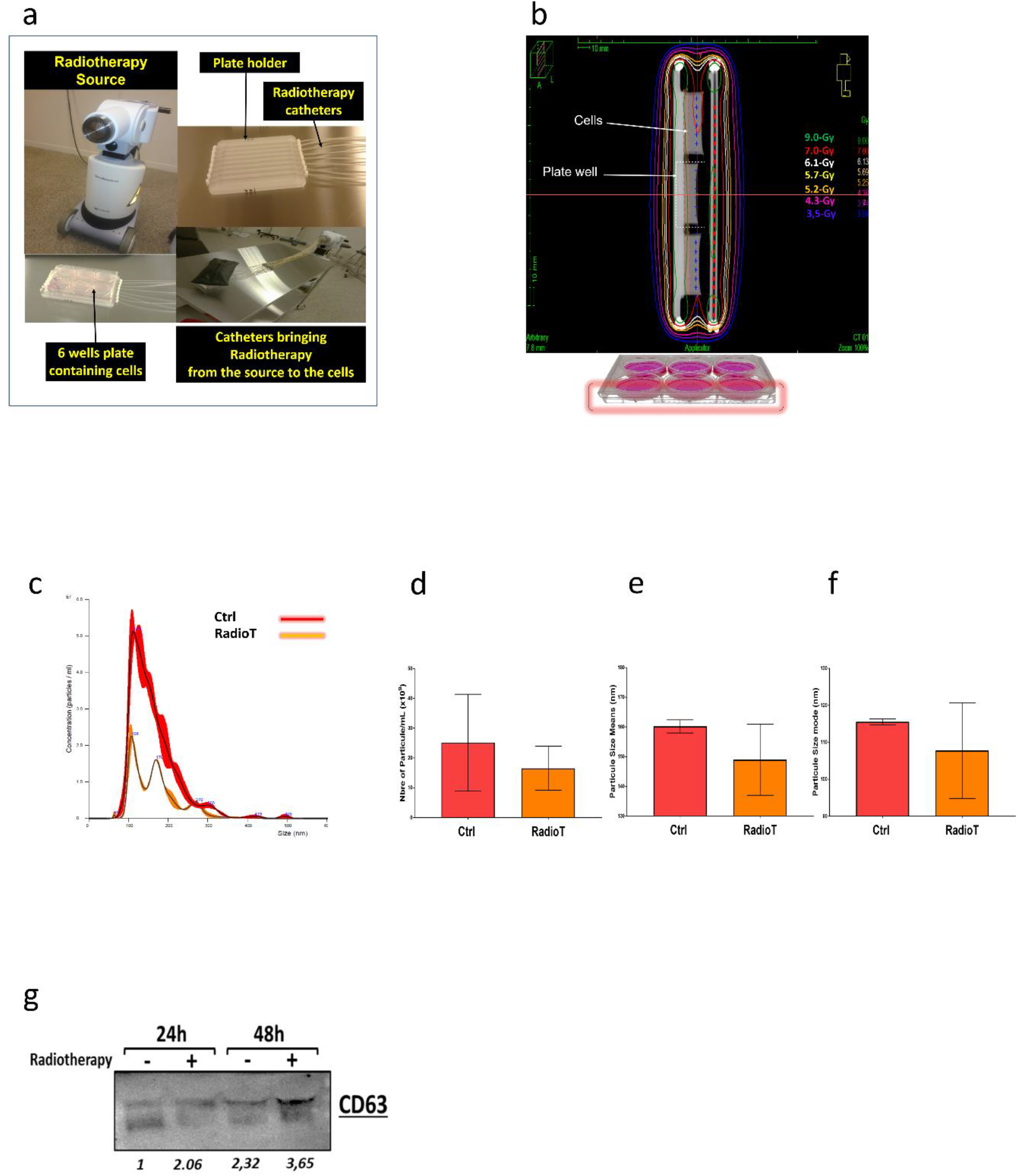
EVs production and sizes from U87-MG cell culture after radiotherapy exposure. (a) The 6 well plates contained U87-MG cells exposed to ionizing radiation. Radiotherapy carried by 8 catheters to the plate holder device, which bears the 6 well plate. All wells received during 20 min a measured dose of 7 Gy. (b) Sagittal cross-section of the 6 well plate, and dosimetry distribution isodoses as decreasing dose layers within the plate from the center to the surface of the plate, isodose in green: 9 Gy, in red: 7 Gy recovering whole cells volume. Then decreasing dosimetry showed in white: 6.13 Gy, yellow: 5.69 Gy, orange: 5.25 Gy, pink: 4.38 Gy, purple: 3.94 Gy and blue: 3.50 Gy. (c,d,e,f): NTA analysis of EVs isolated from 3 mL U87-MG cell supernatants, after 48h culture following radiotherapy exposition (RadioT, orange histograms), or unexposed control cells (Ctrl, red histograms) (fig 1 c). Example of size distribution determined by NanoSight in Ctrl and RadioT conditions. NTA analysis showing the EVs-particle number (fig 1 d), their size means (fig 1 e) and sizes modes (fig 1 f). Histograms represent at least 3 different samples analyzed at least in n=5. (g) Expression of CD63 marker by western-blot, in isolated extracellular vesicle collected and isolated from U87-MG supernatant, 24h and 48h after radiotherapy exposure compared to untreated EVs collected from untreated cells. Numbers indicate the ratio of the western blot bands relative densities values.

### Extracellular vesicles isolation

EVs were isolated from plasma patients and cell line supernatants. Cell lines were cultured at 37°C in a humidified 5% CO2 atmosphere in complete medium (MEM with 5% EVs-free FCS, previously centrifuged for 16 h at 120,000 g to remove bovine exosomes using 40.1Ti rotor (L-80XP; Beckman Coulter^®^, Brea, CA, USA).). Cell supernatants were harvested after 48 h in culture to purify EVs by differential centrifugation as previously described [9].

First, supernatants were centrifuged 10 min at 300 g to remove remaining cells, then 30 min at 16,500 g to remove large extracellular vesicles and other apoptotic vesicles. Next, small EVs were pelleted by ultracentrifugation at 120,000 g for 2 h at 4°C, using 70.1Ti rotor (L-80XP; Beckman Coulter^®^, Brea, CA, USA). EV pellets were wash by PBS and finally, re-centrifuged again at 120,000 g for 2h. Finally, PBS was removed, the size and number of EVs were evaluated using Nanoparticle Tracking Analysis (NTA), NS3000 (Malvern^®^). For further analysis, isolated EVs were suspended in different buffers, depending on the analysis to be carried out (in RIPA lysis buffer for western blot analysis, in RLT buffer, for miRNAs analysis and in PBS for NTA analysis.

### Extracellular vesicles characterization

#### 1) Nanoparticle tracking analysis (NTA)

Vesicle suspensions with concentrations between 1 ×10^7^/ml and 1×10^9^/ml were examined using a Nanosight NS300 (NanoSight Ltd^®^., Amesbury, UK) equipped with a 405 nm laser to determine the size and quantity of particles isolated. Five videos of 60-sec duration was taken for each sample, with a frame rate of 50 frames/sec, and particle movement was analyzed using NTA software (version 3.3; NanoSight^®^ Ltd.).

#### 2) Western blotting analysis

Cells and EVs were lysed in RIPA lysis buffer [50 mM Tris-Cl (pH 8.0), 150 mM NaCl, 1% NP-40, 0.5% sodium Deoxycholate, 0.1% SDS, 100μg/ml phenylmethylsulphonyl fluoride, 0.5 μg/ml leupeptin, and 1 μg/ml aprotinin]. Protein concentrations were determined using a Micro BCA™ protein assay kit (Pierce Biotechnology). Equal amounts of protein were separated by sodium dodecyl sulfate polyacrylamide gel electrophoresis and transferred to PVDF membranes. Membranes were blocked with 5% skim milk in TBS containing 0.1% Tween-20 for 2 h at room temperature and incubated with the appropriate primary and secondary antibodies. Detection of EVs markers was performed with antibodies against CD63 (Abcam, 1/1000) and CD9 (Sigma-Aldrich, 1/1000).

### Radiotherapy treatment of cells

First, prototype device based on the principle of high dose rate (HDR) brachytherapy was developed to expose cell lines to ionizing radiations. The prototype consists in a plexiglas support used as holder to cell lines containing plate. Eight radiotherapy catheters were used to bring ionizing radiation from the source to the Plexiglas support (Figure 1, a). The catheters are connected to an Iridium 192 containing-source projector. This experimental procedure has received agreement N°T870297 from the French nuclear security agency. The targeted volume was measured following dosimetry determined by exposition of six wells-plate to radiotherapy. The radiotherapy levels were measured through the plate volume.

The planning targeted volume corresponding to theoretical cell volume delimited by 7Gy isodose (red line) as observed on plate sagittal sections. Measures showed that the whole volume of U87-MG exposed cells are receiving 7 Gy radiotherapy (Figure 1, b, red layer).

Irradiated cells were then cultured for 24 and 48 hours. The isolation of EVs were carried out from cell supernatants in each condition and their production was evaluated by NTA.

### Collection of plasma from glioblastoma patients

Plasma samples from 4 confirmed glioblastoma patients were collected at four time points during the radiotherapy period noted R0 (before starting radiotherapy), R20 (two weeks after R0, patients receiving 20 Gy), R40 (four weeks after R0, patients receiving 40 Gy) and R60 (six weeks after R0, the end of radiotherapy, patients receiving final dose of 60 Gy).

All patients were informed signed consent and approval forms. The protocol was validated by the ethics committee of Limoges University Hospital, under N°141-2014-08. Plasma samples were stored at −80°C until EVs isolation.

### Total RNA extraction

The miRNAs were extracted using a Trizol extraction method. Cells and EVs were suspended in Trizol Lysis Reagent. Subsequently, high-grade chloroform was added for phase separation and 100% isopropanol for RNA precipitation. Finally, RNA was suspended in 15 to 30 μl RNase-free water after being washed twice in 75% ethanol. Total RNA concentrations were determined using a Nanodrop ND-2000 Spectrophotometer (Labtech, United Kingdom).

### Reverse transcription

Reverse transcription was performed at least on 100 ng of extracted total RNAs, using the TaqMan^®^ MicroRNA Reverse Transcription Kit (Thermo Fisher, USA) following to the manufacturer recommendations and protocols. This reverse transcription method uses a mix of looped reverse transcription primers. Two sets of primers were used: Megaplex RT Primers Human pool A and B (Thermo Fisher, USA). Together, these two pools allow the reverse transcription of 754 different miRNAs.

### Open Array analysis

miRNAs expressions were analyzed using TaqMan Open Array Human MicroRNA Panel plates, containing 754 well-characterized human miRNA sequences from the Sanger miRBase v14. (Thermo Fisher, United States) according manufacturer recommendations and protocols. Pre-amplified cDNA (30 ng) was used for the PCR, and the amplification was done using TaqMan Open Array Real Time PCR 2X Master Mix. The 384-well plate was loaded on the the TaqMan Open Array Human MicroRNA Panel plates (Thermo Fisher, USA) and analyzed using QuantStudio™ 12K Flex (Thermo Fisher, USA). Results were analyzed using Applied QuantStudio Software (Thermo Fisher, USA), and analyzed using 2-ΔΔCt method.

### Statistical analysis

Statistical analysis were performed using R environment (version 3.6.1)[10]. The NormFinder algorithm was used to determine the optimal normalizer miRNA among expressed miRNA [11]. Subsequently, after normalization of the data, the mean of control and radiotreated (RT) samples was used to calculate fold changes (RT control ratio). The significance of differences in means between log2-transformed fold changes was tested using a 2-tailed Welch test after confirming normality by Shapiro test. Significance was set at *p* < 0.05. Exploratory analysis were conducted with principal component analysis (PCA) using FactoMineR package and visualized using factoextra R package [12,13]. Hierarchical clustering analysis were performed with the Complex Heatmap package [14] using Spearman’s correlation coefficient to measure the dissimilarity between each pair of observations and average method to cluster agglomeration.

## Results

### Characterization of EVs from supernatant of U87-MG cells

Following cell irradiation by the Iridium 192 containing-source projector, the extracellular vesicles were isolated from the supernatant of U87-MG cell through a differential ultracentrifugation protocol [9] and were characterized by NTA (Figure 1c). NTA analysis revealed the particles number was similar between the EVs released by control and radiotherapy exposed cells (Figure 1, d). Furthermore, it showed no significant differences in the size means neither modes of the EVs whatever experimental conditions (Fig 1 e, f). Protein analysis on isolated EVs by western blot, showed increase of the expression of CD63 following exposure. This expression was markedly increased after 48-h culture, confirming the increase of EVs production is time dependent (Figure 1, g). Hence, this last time point was chosen to harvest cells and their supernatants. Thereafter, supernatant EVs harvested after 48-h culture were isolated by differential ultracentrifugations. After total RNA purification and extraction from EVs and cells, miRNAs expression were analyzed by Open Array Human miRNAs panel.

### miRNAs expression profile of EVs isolated from cell supernatants before and after irradiation

Analysis of the expression profile of miRNAs in cells was done on 754 human miRNAs screened by the human open array miRNAs panel in U87-MG cells and in their secreted EVs. Less than 10% (9.54 %) of analyzed miRNAs were detected in U87-MG cells, representing 72 miRNAs both expressed before and after irradiation (Figure 2, a, left part), whereas, only 35 miRNAs (about 4.64 %) are detected in the extracellular vesicles isolated from U87-MG cells whatever irradiation or not conditions (Figure 2, a, right part). Interestingly, miRNAs expressed in GBM cells were not detected in their derived-EVs, except for 3 miRNAs.

**Figure 2:**
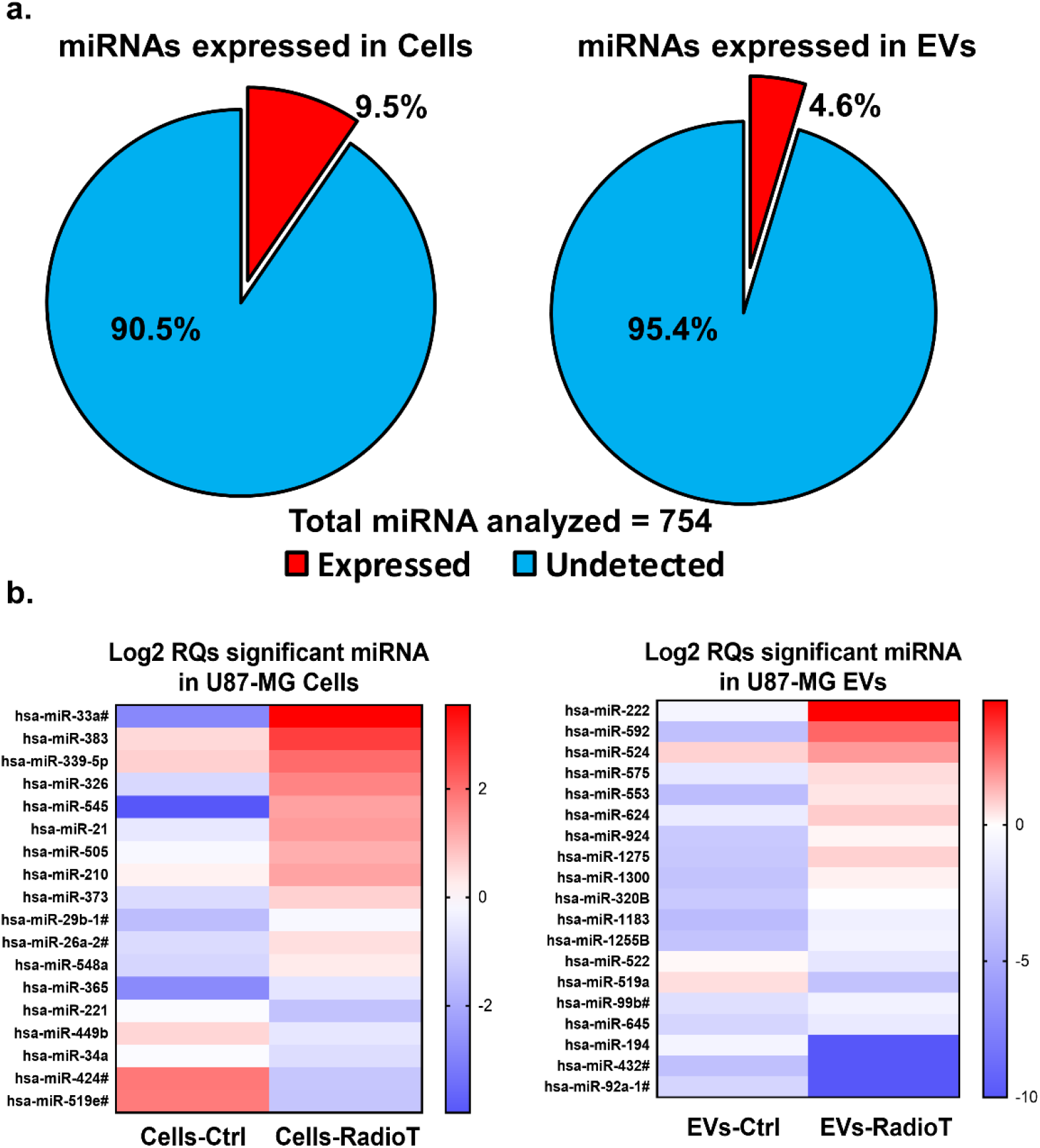
Distribution of miRs differential expression after radiotherapy exposure. (a) Amounts of the detected and expressed miRNA (Red) from whole panel (754 miRNAs) analyzed by Open array on U87-MG cells (left diagram) and on EVs (right diagram) before and after irradiation exposure. (b) Distribution of miRNAs detected in U87-MG (Blue) cells and their produced EVs (Yellow), after 48h culture post radiotherapy exposure. (c) Representative Heat map of significant expressed variation of miRNAs in U87-MG Cells and their isolated EVs before (Ctrl) and after radiotherapy (RadioT) exposure. The variation of miRNAs are represented as Log2, with significant cutoffs of p-value < 0.05, done with T-test with Welch’s correction.

In order to detect a differential expression of miRNAs before and after irradiation, a comparative analysis of miRNAs expression between cells and EVs was performed and represented by heatmap. Indeed, a differential miRNA expression between non-irradiated control U87-MG cells (Cells-Ctrl condition) and cells irradiated with 7 Gy (Cell-RadioT condition) was evidenced (Figure 2, c, left part). Similarly, the expression profile of miRNAs analyzed in the secreted-EVs from treated cells (EVs-RadioT condition) was compared to the expression profile of miRNAs in EVs-isolated from untreated control cells (EVs-Ctrl condition) (Figure 2, c, right part).

Results showed a significant variation of miRNAs expression induced by irradiation marked by the overexpression of 12 miRNAs and a decreased expression of 6 miRNAs. (Figure.2, c, left part). In parallel, 12 miRNAs were significantly overexpressed, whereas 9 miRNAs were decreased in extracellular vesicles after irradiation (Figure 2, c, right part). Surprisingly, none of these miRNAs expression profiles induced by irradiation, shares a common significant expression of miRNAs between U 87-MG cells and their secreted EVs (Figure 2, c).

According to the variations of miRNAs expression in EVs from of U-87MG cells after irradiation, we analyzed EVs miRNA contents from blood samples of 4 patients with GBM during the course of radiotherapy treatment.

### Characterization and variation of EVs isolated from GBM patients plasma during radiotherapy treatment

A sampling process was defined to collect glioblastoma patient’s plasma during radiotherapy protocol. Four blood samples were collected at different time points at R0 (before starting radiotherapy), R20 (at 2 weeks after the start of radiotherapy, patients received 20 Gy), R40 (four weeks after starting radiotherapy, patients received 40 Gy) and R60 (at six weeks after starting treatments, patients received 60 Gy, corresponding to the end of radiotherapy) (Figure 3, a).

**Figure 3:**
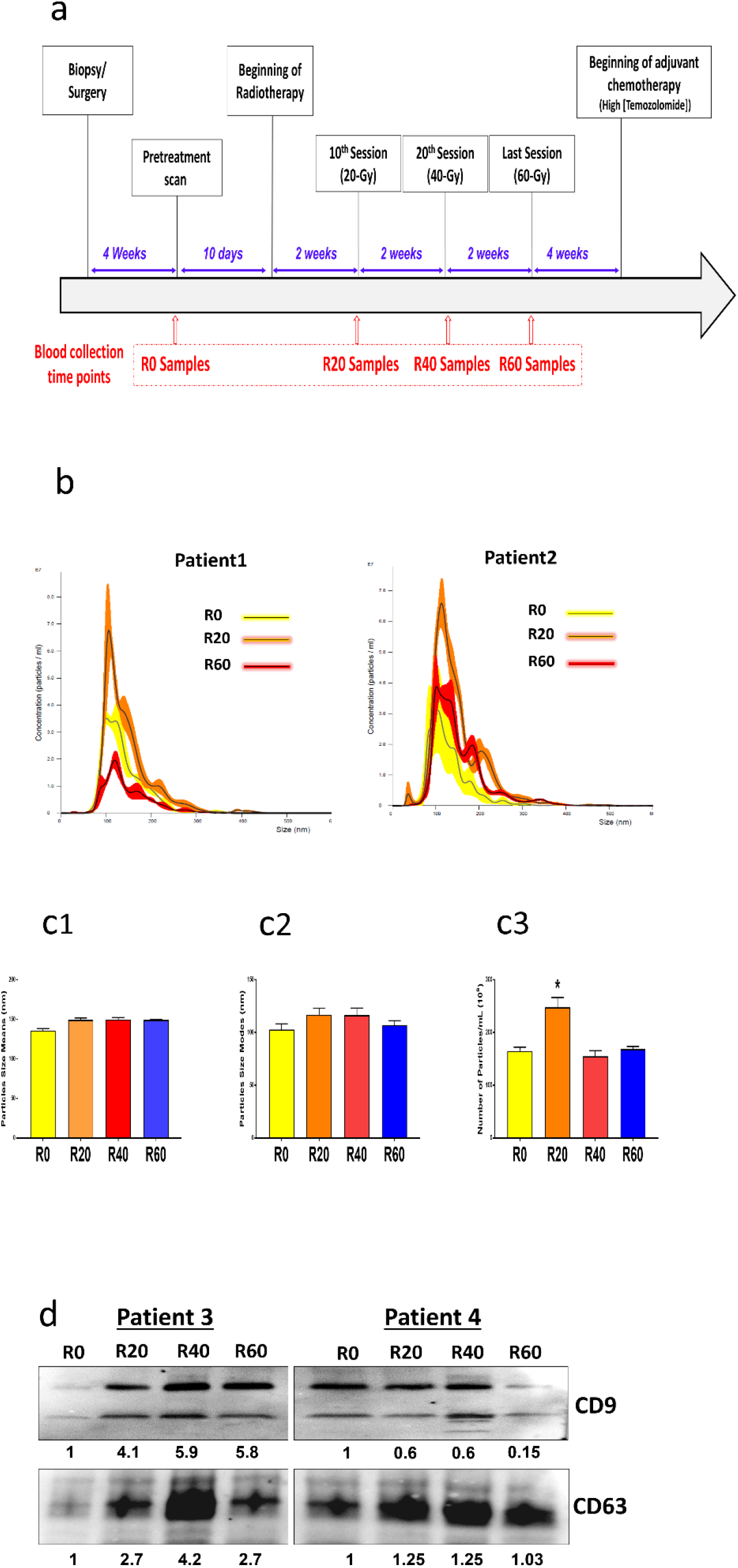
Patients’ radiotherapy induced an increase of EVs production in R20 plasma samples. (a) Kinetic of Evs in plasma from glioblastoma patients during radiotherapy treatment following the called Stupp protocol (Stupp et al. 2005). R0 corresponded to samples collected before starting radiotherapy protocol, R20, R40 and R 60 samples were collected 2 weeks, 4 weeks and 6 weeks respectively after the beginning of radiotherapy, with a cumulative irradiation of 20 Gy, 40 Gy and 60 Gy, respectively. (b) NTA analysis of EVs concentration (particles/mL) isolated from 1 mL of 2 different patients’ plasma at R0 (yellow), R20 (orange) and R60 (red) time-points. (c) A representative NTA analysis of plasma Evs distribution evaluated by NanoSight in R0, R20 and R60 samples showing their size means (C1), sizes modes (C2) and EVs-particle numbers (C3). * (*P* value < 0.05). (d) Representative western-blot analyzes of CD63 and CD9 markers expression in isolated EVS collected and isolated from R0, R20, R40 and R60 plasma samples. Isolated EVs were resuspended in lysis buffer for protein extraction. Proteins (50 μg) were loaded and analyzed for each sample.

Extracellular vesicles isolated from plasma samples, were then isolated and characterized. NTA analysis showed picks with narrow profiles between EVs samples isolated from different conditions R0, R20 and R60 (Figure 3, b). Particle size means were 135 to 150 nm (Figure 3, c1) with modal diameter of 102 to 118 nm (Figure 3,c2). The average of EVs particles number was 150 to 250 x 10^9^, isolated from 1 mL of plasma in each condition (Figure 3, c3). Interestingly, particle number significantly increased at R20 (Figure 3, c3) by comparison with before (R0) and the end of radiotherapy (R40 and R60). However, no significant difference was observed in the size of isolated EVs (Figure 3, c2, c3). The expression of two markers of exosomes, CD9 and CD63, analyzed by western-blot confirmed the increase of the EVs secretion at R20 and R40 compared to the control condition R0. (Figure 3, d) with a decreasing tendency of EVs markers expression at R60 (Figure 3, d).

For further analysis of EVs isolated from patients, we analyzed the differential expression of miRNA content after radiotherapy exposure at R20 and R60 conditions in comparison with the condition R0, before starting radiotherapy.

### miRNAs expression profile in EVs isolated from plasma patients

As previously shown in EVs isolated from the U87-MG cell line, Open Array analysis was done on 754 miRNAs in EVs isolated from patients’ plasma at different level of radiotherapy protocol corresponding to distinct doses R0 (0 Gy), R20 (20 Gy) and R60 (60 Gy). This study detected 14.7% miRNAs expressed in patients’ EVs corresponding to 111 different miRNAs (Figure 4a). The principal component analysis has revealed three patterns of expression for the analysis done on plasma-derived EVs from four GBM patients, depending on the radiotherapy treatment stage (Figure 4 b). Irrespective of the considered patient, miRNAs expressed in R0 condition form an independent group (in red), which is separated from the treated conditions R20 (in green), demonstrating that the treatment R20 generates a miRNAs profile different from the R0 control one. However, the observed profiles of R60 group (in blue) show that do not have an expression profile overlapping with R20 group nor with the control group R0 (Figure 4, b).

**Figure 4:**
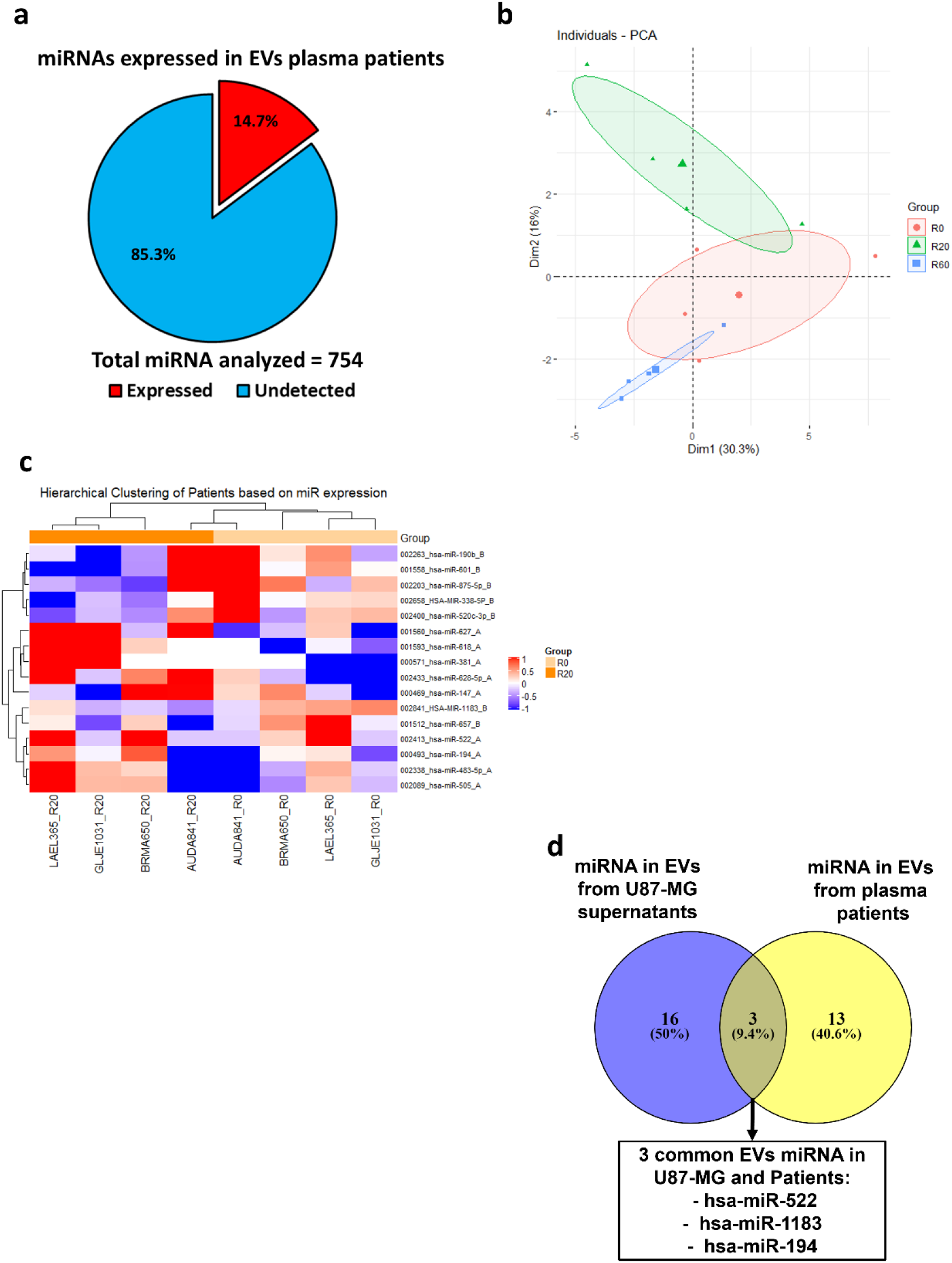
Differential expression of miRs in plasma EVs from patients during radiotherapy. (a) Detection of expressed miRNA (Red) in isolated EVs plasma from 4 GBM patients analyzed in whole panel (754 miRNAs) by Open Array. (b) Principal component analysis (PCA) of expression patterns of miRNAs in EVs isolated from 4 plasma patients before radiotherapy exposure (R0) and after 20 Gy (R20) or 60 Gy (R60) radiotherapy treatment. (c) Two-way hierarchical clustering of significant variation of expressed miRNAs in EVs isolated from plasma of GBM patients at R20 compared to R0 condition. The variation of miRNAs are represented as Log2, with significant cutoffs of p-value <0.05, done with T-test with Welch’s correction. (d) Venn diagram presenting the comparison of miRNAs differential expression in EVs from U87-MG cell supernatants (before irradiation and after Cell irradiation) and plasma patients at R0 and after 20 Gy irradiation). Three miRNA are commonly expressed in EVs following irradiation conditions (U-87MG cells and patients at 20 Gy).

Two-way hierarchical clustering of expressed miRNAs in EVs isolated from GBM patients’ plasma showed that 16 miRNA have a significantly different expression between the R0 control and the R20 radiotherapy conditions, providing an expression pattern for each analyzed condition (Fig 4, c and Table 1a). Thus, this observation confirmed the patterns highlighted by the PCA showing the differential expression of miRNAs in EVs before radiotherapy exposure R0 and after 20 Gy radiotherapy treatment (R20 condition).. The Venn diagram shows that only 3 miRNAs (hsa-miR-522, hsa-miR-1183 and hsa miR-194) are common to EVs from U87-MG cells after irradiation and EVs from irradiated patients with a total dose of 20 Gy (Figure 4d). In addition, miR522 is the single to be overexpressed in EVs when patients are exposed to different doses of radiation therapies (Table 1a and Fig. S1b) In addition, it is the only one of the 3 miRs whose overexpression tends to be associated with poorer survival (p=0.061) (Fig.S2a and 2b). Since these 3 miRNAs are isolated from EVs and specifically expressed following irradiation, their expression could be closely linked to cell changes occurring after radiotherapy. Overall, these results suggest that these miRNAs could represent a molecular profile to analyze cellular modifications after radiotherapy.

**Table 1.**
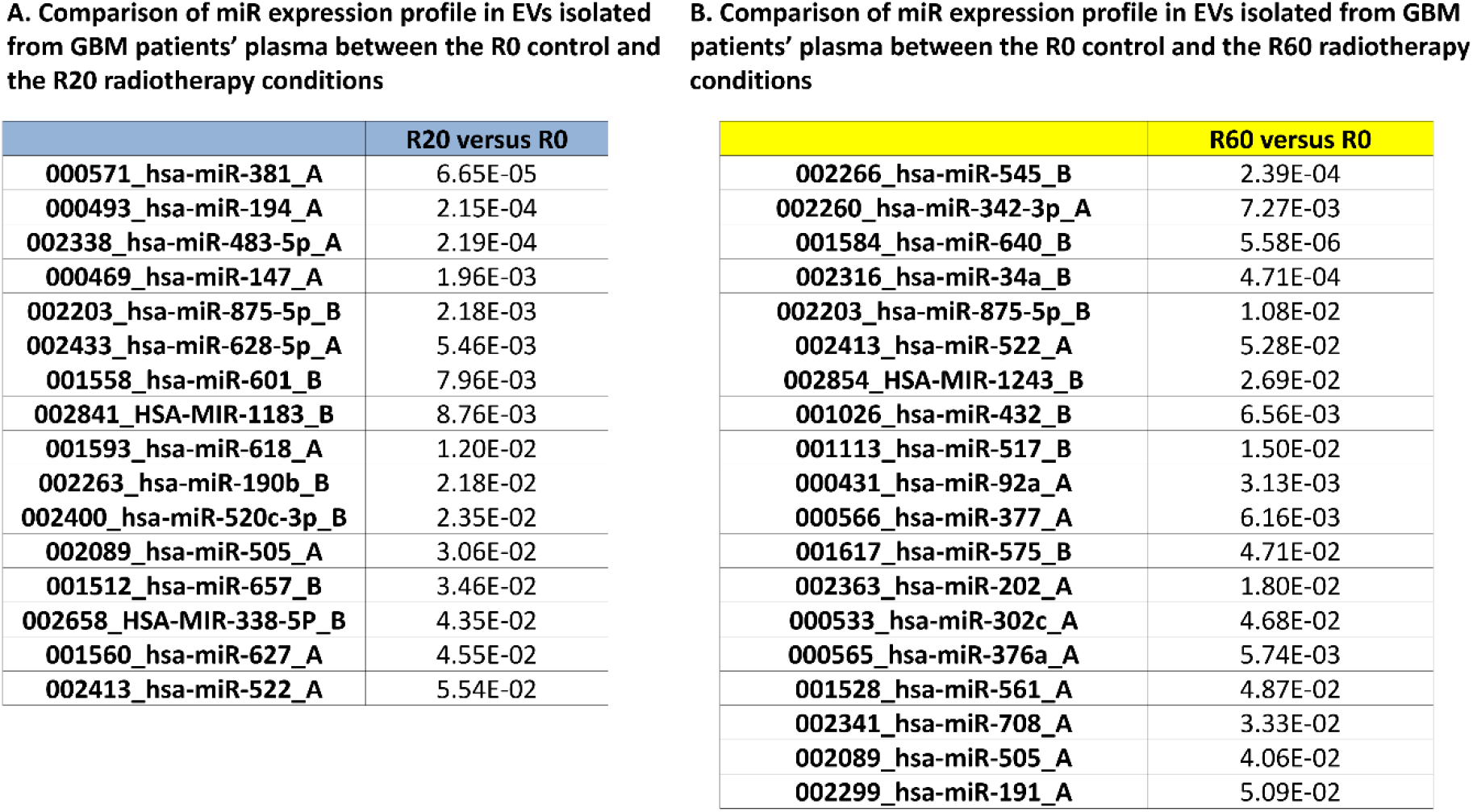
P values correspond to t test performed for mean miR expression comparison between radiotherapy conditions (R20 for A and R60 for B) and R0 control after log2 transformation of RQ values calculated by ΔΔCt method. A. Comparison of miR expression profile in EVs isolated from GBM patients’ plasma between the R0 control and the R20 radiotherapy conditions B. Comparison of miR expression profile in EVs isolated from GBM patients’ plasma between the R0 control and the R60 radiotherapy conditions

|  | **R20 versus R0** |
| --- | --- |
| 000571_hsa-miR-381_A | 6.65E-05 |
| 000493_hsa-miR-194_A | 2.15E-04 |
| 002338_hsa-miR-483-5p_A | 2.19E-04 |
| 000469_hsa-miR-147_A | 1.96E-03 |
| 002203_hsa-miR-875-5p_B | 2.18E-03 |
| 002433_hsa-miR-628-5p_A | 5.46E-03 |
| 001558_hsa-miR-601_B | 7.96E-03 |
| 002841_HSA-MIR-1183_B | 8.76E-03 |
| 001593_hsa-miR-618_A | 1.20E-02 |
| 002263_hsa-miR-190b_B | 2.18E-02 |
| 002400_hsa-miR-520c-3p_B | 2.35E-02 |
| 002089_hsa-miR-505_A | 3.06E-02 |
| 001512_hsa-miR-657_B | 3.46E-02 |
| 002658_HSA-MIR-338-5P_B | 4.35E-02 |
| 001560_hsa-miR-627_A | 4.55E-02 |
| 002413_hsa-miR-522_A | 5.54E-02 |

|  | **R60 versus R0** |
| --- | --- |
| 002266_hsa-miR-545_B | 2.39E-04 |
| 002260_hsa-miR-342-3p_A | 7.27E-03 |
| 001584_hsa-miR-640_B | 5.58E-06 |
| 002316_hsa-miR-34a_B | 4.71E-04 |
| 002203_hsa-miR-875-5p_B | 1.08E-02 |
| 002413_hsa-miR-522_A | 5.28E-02 |
| 002854_HSA-MIR-1243_B | 2.69E-02 |
| 001026_hsa-miR-432_B | 6.56E-03 |
| 001113_hsa-miR-517_B | 1.50E-02 |
| 000431_hsa-miR-92a_A | 3.13E-03 |
| 000566_hsa-miR-377_A | 6.16E-03 |
| 001617_hsa-miR-575_B | 4.71E-02 |
| 002363_hsa-miR-202_A | 1.80E-02 |
| 000533_hsa-miR-302c_A | 4.68E-02 |
| 000565_hsa-miR-376a_A | 5.74E-03 |
| 001528_hsa-miR-561_A | 4.87E-02 |
| 002341_hsa-miR-708_A | 3.33E-02 |
| 002089_hsa-miR-505_A | 4.06E-02 |
| 002299_hsa-miR-191_A | 5.09E-02 |

The analysis of two-way hierarchical clustering of expressed miRNAs in EVs isolated from GBM patients’ plasma at R0 compared to R20 or R60 also revealed that a couple of miRs are significantly expressed before or after irradiation (Table 1a and.1b). We mainly focused on miR-381 which is markedly up-regulated after a dose of 20Gy in radiotherapy (Fig. 4C). Unlike, its expression is not significantly increased in EVs from patients that have undergone radiotherapy at doses 60 Gy (Fig. S1a and Table 1b). Nevertheless, miR-381 overexpression is associated with lower overall survival unveiling the increase of its expression upon radiotherapy could afford a marker of poor prognosis. (Fig. S2).

### Identification of miRNA targeted mRNAs derived from EVs from plasma of patients upon radiotherapy

To determine presumed targets of four deregulated miRNA, we chose to use six online prediction tools using algorithm default parameters. Six distinct genes list were obtained from each prediction tool (Table S1a). The number of target RNAs for a given miRNA according to the different online prediction tools or for the 4 miRNAs obtained from the same prediction tool were very heterogeneous and directly depended on the chosen online prediction tool (Table S1a). The miRNA targets from each list were cross-referenced to characterize the putative common targets of the 4 miRNA selected in our study. The greater the number of lists in which the target is expressed, the more it is a target of interest. A single gene, CADM1 is present in 4 different lists (Table S1b). Thus, CADM1 can be considered a specific target for these 4 miRNAs. Thereby, it might afford a sensitive marker to radiations.

## Discussion

Tumors have a remarkable ability to control and modulate their microenvironment to their own advantage. These properties contribute to resistance mechanisms by promoting the communication between cancer cells and their surroundings. EVs released by cells within the tumor microenvironment in response to anti-cancer therapies have also been shown to mediate resistance [15]. However, the function of exosomes in resistance mechanisms is still unclear. Although the tumors harbored very homogeneous miRNAs expression profiles in general, we could identify several miRNAs that were differentially expressed in cells and exosomes and 3 miRNAs common between exosomes derived from cells and patients. We confirm as Arscott et al. that cellular irradiation increases exosome secretion as the number of exosomes is markedly increased after ionizing radiations [8]. Furthermore, we found that irradiation significantly increased the total protein content of the released EVs. Nevertheless, Arscott observed few changes in their miRNAs composition except for miRNAs implicated in cell movement. In contrast to Arscott’s results, irradiation induces changes of miRNAs expression in radiation-derived exosomes. Indeed, 19 miRNAs are differentially expressed in cell derived-exosomes following radiotherapy. This discrepancy can be due to high irradiation dose in our experiments (7 Gy) while Arscott performed irradiation at 4 Gy. This result suggest the irradiation doses might modify the miRNA content in exosomes. These data are supported by previous findings in breast cancer cells showing that when cells damage are increased in a dose-dependent manner, exosomes secretion and activity of exosome secretion pathway were enhanced [16].

Our study shows that few miRNAs contained in exosomes are similar between the GBM cell line and the plasma from glioblastoma patients. This could be due to the release of exosomes of cells from the microenvironment such as cancer-associated fibroblasts or mesenchymal stem cells (MSCs) that secrete miRNAs into exosomes in order to promote cell migration or invasion in different cancers [17]. Similar mechanisms can be observed in MSCs re-programmed by tumor-derived EVs that in turn produce their own exosomes. mRNA and miRNA species as well as molecular signals are delivered back not only to tumor cells to enhance their growth, but also to fibroblasts, endothelial cells and immune cells in the microenvironment to promote tumor cells growth, angiogenesis and immune escape [18]. Finally, only 3 miRNAs (miR-1183, miR-194, miR-522) were common to cell derived-exosomes and patients derived-exosomes. However, miRNAs from cells and exosomes are differentially expressed and regulated. Thus, in exosomes derived from GBM cells, miR-194 and miR-522 are underexpressed following ionizing radiations whereas miR-1183 is overexpressed. In contrast, the expression level of same miRNAs are totally opposite in patients derived-exosomes in which miR-194 and miR-522 are upregulated, unlike miR-1183 is downregulated The significant changes in miR profiles *in vitro* and in patients could depend on the influence of tissue microenvironment, which is absent *in vitro*. Changes of cell culture and environment can influence the cell proliferation and select cell subpopulation carrying specific patterns of miRNAs. In this context, culture conditions could modify the genotype and phenotype of neoplastic cells and influence their miR profile [19]. For this reason, in our study we chose to analyze mainly the changes in the expression levels of the 3 common miR in patient-derived exosomes. Among these 3 common miRNAs, miR-194 is significantly overexpressed upon ionizing radiation. As previously described, its expression significantly increased in patients responding to preoperative chemoradiotherapy in advanced rectal cancer [20]. Indeed, it is likely that miR-194 expression can depend on radiation doses (7 Gy in our experiments), since it is already known to be regulated at high absorbed doses used for the treatment of neuroendocrine cancers [21]. The expression of miR-194 is clearly correlated with a tumor suppressor function in various cancers including hepatocarcinoma in which it regulates PTBP1 expression involved in tumor proliferation and migration, but also in glioblastoma [22,23]. Altogether, these results suggests miR-194 upregulation could afford a predictive biomarker of response to radiotherapy. Nevertheless, the exact function of miR-194 in tumor released-exosomes could lead to controversy. Indeed, it exerts angiogenic functions through proliferation of pulmonary microvascular endothelial cells, in murine and human colon cancer cell lines and activates the astrocyte-endothelial cell transition [24,25]. Consequently, it might be involved in tumor angiogenesis and could participate to gliomagenesis and tumor progression. The expression of miR-522 in exosomes undergoes the same changes after irradiation as miR-194 [24]. Its expression is associated with tumor progression and growth in several cancers [26,27]. In glioblastoma, miR-522 is also known to promote tumor cell proliferation by downregulating a Ser/Thr protein phosphatase (PHLPP1) involved in maintaining suppression of cell survival to prevent oncogenesis [28]. Its expression significantly increases in tumor cells and tissues, in contrast to adjacent non-tumor brain tissues and healthy astrocytes. However, miR-522 expression is not restricted to tumor tissue alone but is also detected in circulating EVs in non-small cell lung cancer [29]. Indeed, the transfer of miR-522 via exosomes represents a specific mechanism of propagation and transfer of tyrosine kinase inhibitors (TKI) resistance to neighboring cells that contributes to the increase of miR-522 in EGFR inhibitor-resistant cells.

Our results confirmed miR-522 up-regulation in EVs derived from glioblastoma patients. In this cancer, the transfer of miR-522 to exosomes after irradiation had never been observed before. For the first time, our results demonstrate that miR-522 expression is based on ionizing radiations as confirmed by the higher expression of miR-522 after 20 Gy irradiation in glioblastoma. This upregulation and specific transfer into exosomes persists at 60 Gy suggesting that its expression in exosomes is closely related to the response to ionizing radiations. Although its expression in 562 glioblastoma patients before therapy, analyzed with TCGA database, shows that miR-522 is not closely associated with overall survival (p= 0.061, Fig. S2), its increase in the EVs loadings after radiotherapy suggests that miR-522 could be a potential marker of poor prognostic for response to radiotherapy. In comparison with the role of miR-522 in the transmission of TKI resistance, further experiments will be required in order to analyze the exact function of miR-522 transfer in recipient cells and its involvement in radioresistance mechanisms [29]. The expression of miR-1183 is in opposition to both the two other miRs. This result corresponds to the first observation of this miR in extracellular vesicles. Moreover, its expression in exosomes is clearly associated with radiotherapy as confirmed by the decreasing of its expression to 20 Gy, which continues to 60 Gy. Since miR-1183 belongs to a subset of 13 miRNAs that are correlated with the complete pathological response after neo-adjuvant chemo-radiotherapy in patients with rectal cancer, it could be considered as a potential marker to predict a response to treatment [30]. In glioblastoma, the down-regulation of miR-1183 in exosomes released from irradiated glioblastoma cells or tumor patients could suggest that this miR might not play its tumor suppressor role on recipient cells. Since the process of tumor irradiation could be assimilated to cellular stress conditions with microenvironmental adaptations, it involves the release of specific miRNAs. As miR-1183 was not detected in the TCGA patients database, we are not able to establish the correlation between its expression and clinical data. However, we can hypothesize that the decrease of miR-1183 in circulating exosomes might reflect the level of miR expression in tumors. Thus, the miR-1183 down-regulation associated with miR-522 and miR-194 up-regulation might represent a new reliable and specific miR profile linked to radiotherapy response. In addition, an other miRNAs, miR-381, not expressed in U87-MG cells, were also selected since they are significantly up-regulated upon 20Gy irradiation and could be useful to complete miRNAs profile in response to radiotherapy (Fig.S1). The miR-381 expression is variable in cancers, but is generally correlated with tumor regression and patient’s survival [31]. However in glioblastoma, it plays an opposite role since miR-381 increase promotes tumor growth [32]. These data are consistent with our preliminary results and are supported by the Kaplan-Meier analysis of the TCGA database that shows that high expression of miR-381 decreases survival in 562 glioblastoma patients (p = 0.01, Fig S2) [33].

Altogether, these results suggest the definition of a new miR profile defined by 4 selected miRNAs (miR-1183, miR-194, miR-522, miR-381). Six online prediction tools using algorithm default parameters allowed characterizing six distinct genes list (Table S1a). A single gene, CADM1 is common in 4 different lists and might be a putative common target for all selected miRNA. CADM1 is considered a tumor suppressor gene which is sensitive to radiotherapy as supported by CADM1 alteration following thorax irradiation [34,35]. Moreover, its expression is negatively regulated in glioblastoma by an other miR, miR-148a [36]., miR-148a was transferred by GBM cells-released EVs for enhancing the proliferation and invasion. Given previous works, we can hypothesize the 4 miRNA characterized after radiotherapy in the present study might be involved in CADM1 downregulation in response to radiotherapy in order to promote tumor progression.

This miRs subset and their single common target could be useful and reliable to detect and monitor miRNAs changes after radiotherapy. Consequently, the establishment of a comparative exosomal microRNA profile at different levels could facilitate the characterization of tumor aggressiveness and assess the risk of radioresistance. Exosomes-content miRNA therefore afford a promising biological marker for monitoring radiotherapy response and potential optimization of glioblastoma treatment. Furthermore, this specific miR signature might have a clinical relevance in order to optimize the therapeutic management of patients in future. Nevertheless, further experiments in a larger patient population with clinical data will be required to define whether this signature might have a prognostic value on radiotherapy response.

## Supplementary Materials

**Figure S1: (a) Differential expression of miRs in plasma EVs from patients after 60Gy irradiation; (b) Common deregulated miRs in EVs isolated from GBM patients between the R0 control and radiotherapy (Venn diagram)**

**Figure S1a:** Two-way hierarchical clustering of significant expressed variation of miRNAs in EVs isolated from plasma of GBM patients at R60 and R0 conditions. The variation of miRNAs are represented as Log2, with significant cutoffs of p-value < 0.05, done with T-test with Welch’s correction.

**Figure S1b:** 3 miRs are significantly deregulated to both conditions R60 and R20.

**Figure S2: Kaplan-Meier analysis for overall survival depending on changes of miR-522, miR-194, miR-381 expression levels.**

Low miR expression (green) and high miR expression (red) were defined by the cutoff determined by cutpoint of survminer package R. Higher miR-381 expression is significantly correlated with the longer survival of patients with Glioblastoma. The expression level of both miRNA, miR-522 and miR-194, is not associated with survival changes in these patients. (*P value < 0.05).

**Table S1: In silico research of mRNA target of 4 selected miRNA deregulated in EVs from plasma patients during radiotherapy.**

## Ethics approval and consent to participate

All samples were used in accordance with French bioethic laws regarding patient information and consent. Patients’plasma were obtained from Limoges University Hospital, following research project approval by the Institutional Review Board (N°141-2014-08).

## Consent for publication

All authors provided consent for publication

## Availability of data and material

All patient samples and data are available and stored in the Tumor Biobank of Limoges University Hospital.

## Author Contributions

Conceptualization, Pierre Clavère, Marie-Odile Jauberteau and Fabrice Lalloué; Formal analysis, Sofiane Saada, Stephanie Durand and Fabrice Lalloué; Investigation, Sofiane Saada, Stephanie Durand, Alexandre Nivet, Barbara Bessette, Axel Boukredine and Amel Rehailia; Methodology, Sofiane Saada, Stephanie Durand, Alexandre Nivet, Barbara Bessette, Amel Rehailia, Pierre Clavère and Elise Deluche; Supervision, Fabrice Lalloué; Validation, Elise Deluche and Fabrice Lalloué; Writing – original draft, Sofiane Saada, Stephanie Durand, Elise Deluche and Fabrice Lalloué; Writing – review & editing, Pierre Clavère, Marie-Odile Jauberteau and Fabrice Lalloué.

