## Supplementary figures and images for "Radiotherapy induces specific miRNA expression profiles in glioblastoma exosomes"

### Figure S1

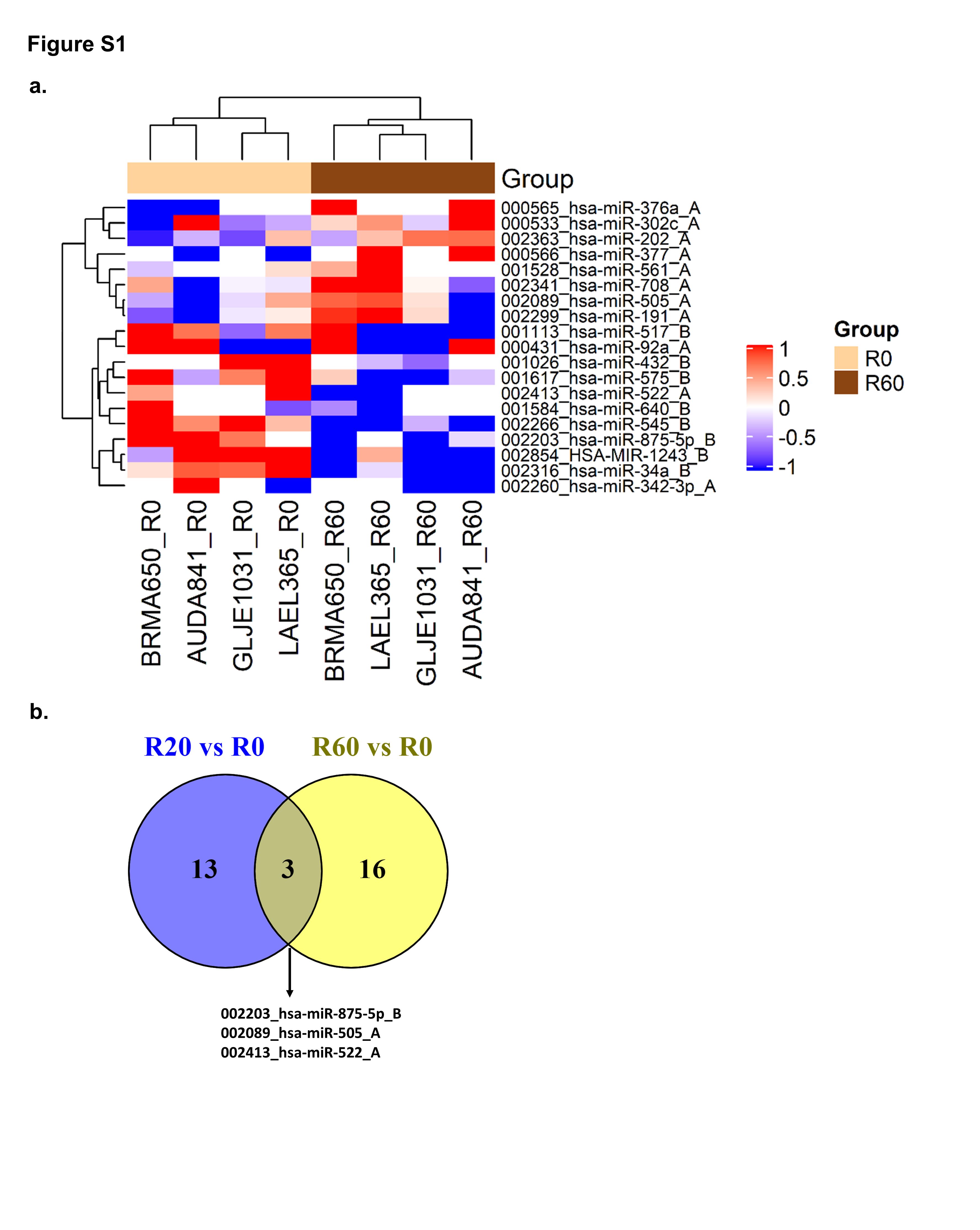

### Figure S2

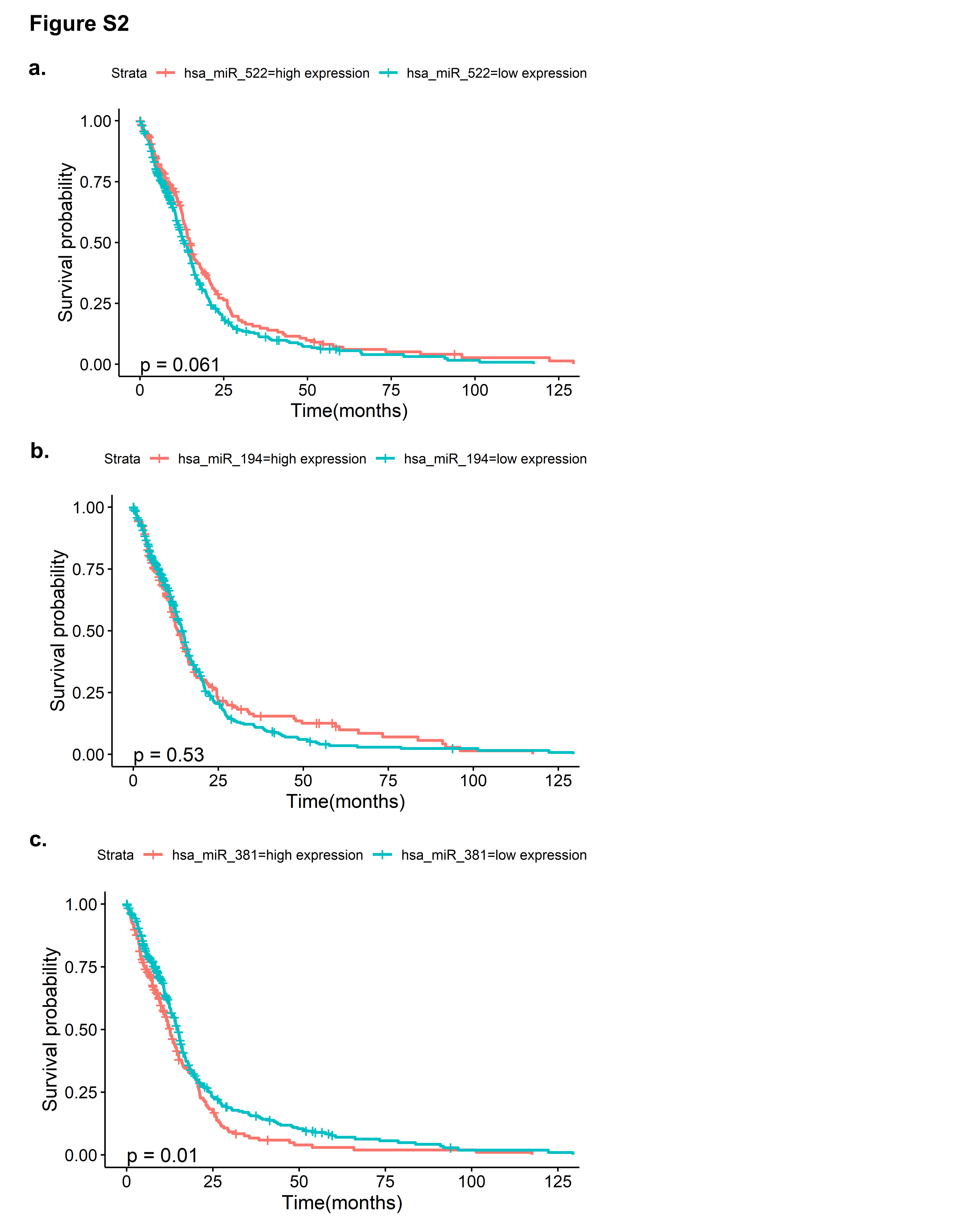
