## Supplementary material for "Radiotherapy induces specific miRNA expression profiles in glioblastoma exosomes": Table S1

**Table S1**. ***In silico* research of mRNA target of 4 selected miRNA deregulated in EVs from plasma patients during radiotherapy.**

**A.** Identification of miRNA targets across six online prediction tools using algorithm default parameters (miRDB v6.0, miRWalk v3.0, TargetScan v.7.2, DIANA-microT-CDS v.5.0, microRNA.org and TargetMiner).

Each list of target of each miRNA were crossing to identify common targets of 4 miRNA.

| **miRNA name (Accession Number)** | **miRDB** | **miRWalk** | **TargetScan** | **DIANA-microT-CDS** | **microRNA.org** | **TargetMiner** |
| --- | --- | --- | --- | --- | --- | --- |
| **hsa-miR-1183 (MIMAT0005828)** | 517 | 1903 | 3539 | 777 | 3248 | 1415 |
| **hsa-mir-522-3p (MIMAT0002868)** | 836 | 940 | 4803 | 2038 | 3849 | 1579 |
| **hsa-miR-194-5p (MIMAT0000460)** | 676 | 672 | 473 | 860 | 2880 | 1351 |
| **hsa-miR-381-3p (MIMAT0000736)** | 1320 | 1348 | 833 | 2293 | 4294 | 1379 |
| **Common targets of 4 miRNA** | **9** | **6** | **35** | **30** | **217** | **219** |

Web access of prediction tools:

miRDB: <http://www.mirdb.org/> ; miRWalk : <http://mirwalk.umm.uni-heidelberg.de/> ; TargetScan : <http://www.targetscan.org/vert_72/> ; DIANA-microT-CDS : <http://diana.imis.athena-innovation.gr/DianaTools/index.php?r=microT_CDS/index> ; microRNA.org : <http://www.microrna.org/microrna/home.do> ; TargetMiner : <https://www.isical.ac.in/~bioinfo_miu/targetminer20.htm>

**B.** Crossing of common targets of four miRNA sorting from six different prediction tools.

| **Number of Prediction Tools** | **Number of Common Targets** | **Target Name** |
| --- | --- | --- |
| 4 | 1 | CADM1 |
| 3 | 6 | UBR3, NEUROD1, NFAT5, ACVR2B, SOCS5, TAOK1 |
| 2 | 35 | SCAMP1, WAPAL, SALL1, PTCH1, GPCPD1, PTPRD, LRP12, RAB11FIP2, FNBP1L, SLC2A13, QKI, KIF1B, SP4, DCUN1D4, NAB1, ST8SIA4, NF1, DACH1, UBE2W, TEAD1, VAPA, PHC3, KLF12, ARHGAP5, EIF4E, DR1, DGKH, FZD3, RSF1, KALRN, CCDC144NL, TNRC6A, TSHZ1, ANO1, GDAP2 |
