## Supplementary materials for "Radiotherapy induces specific miRNA expression profiles in glioblastoma exosomes"

**miR expression and Survival analysis from TCGA data**

The Cancer Genome Atlas (TCGA) data for normalized and batch-corrected miRNA expression of glioblastoma (GBM) cohort were downloaded from the Firehose portal of the Broad Institute (https://gdac.broadinstitute.org/). Clinical informations were retrieved also from Broad GDAC portal and from Genomic Data Commons Data Portal of National Cancer Institute (https://portal.gdc.cancer.gov/projects/TCGA-GBM). TCGA-GBM cohort with miRNA expression profiling consist of 562 patients (219 women and 343 men) with 59 years of median age at initial diagnosis. During follow-up of cohort, 426 patients were deceased.

In a second time, we have selected two subsets of patients based on therapy:

Survival analysis were performed in R environment using survival (version 2.38) and survminer (version 0.4.6) packages. Each cohort were stratified according miRNA expression level. Briefly, the R Maxstat function was used to compute the cutoff yielding to the maximum difference in overall survival between patients with low or high miRNA expression (survminer package). The minimal proportion of observations per stratified group was set to 0.3. Survival curves were obtained using the Kaplan–Meier method (p values indicated in the graphs correspond to log-rank test). Relevant variables associated with overall survival (OS) were examined using univariate. Tests or comparisons were considered significant when p ≤ 0.05.
